# Structural basis for phosphorylation-induced inactivation of the rice ammonium transporter AMT1;1

**DOI:** 10.64898/2026.10.12.758405

**Authors:** Fang Yu, Xiaowei Huang, Shuang Wang, Ning An, Peng Zhang, Jirong Huang, Xue Zhang

## Abstract

Ammonium (NH_4_^+^) uptake and transport in plants are mediated by ammonium transporters (AMTs), which are tightly regulated by phosphorylation to prevent excess ammonium influx and toxic accumulation. However, the mechanism of phosphorylation-mediated inactivation is unique and remains unclear. Here, we report cryo-EM structures of the rice (*Oryza sativa*) ammonium transporter OsAMT1;1 in its wild-type and phosphomimetic variant T452D states. OsAMT1;1 forms a stable homotrimer, with conserved motifs affirming a shared transport mechanism within the AMT/MEP/Rh superfamily. Our structural analysis reveals that T452D phosphomimetic stabilizes an intracellular salt-bridge network and maintains the channel in a closed state, in which the ICL5 loop positions the hydrophobic plug residue Leu360 near the cytoplasmic exit to occlude the channel exit and abolish complementation capacity. Our findings elucidate phosphorylation-induced inactivation mechanism for the plant AMT1 subfamily, providing a structural paradigm for the post-translational regulation of nutrient uptake.

## Introduction

Nitrogen (N) is a primary component of the nucleotides and proteins that are essential for life [1]. Ammonium (NH_4_^+^) and nitrate (NO_3_^‒^), the major inorganic N forms in soil, are crucial for crop growth and yield formation. In aerobic soils, the major form of inorganic N is NO_3_^‒^, but NH_4_^+^ is the predominant N form in flooded wetland or acidic soils [2]. Plants preferentially absorb ammonium since its metabolism requires less energy than processing nitrate [3,4]. The concentration of ammonium nitrogen in soils typically remains below 1 mM [5]. The uptake and transport of ammonium in plant roots are mediated by plasma membrane-localized ammonium transporters (AMTs), which belong to the evolutionarily conserved AMT/MEP/Rh superfamily, a ubiquitous transmembrane protein group identified in diverse organisms spanning bacterial, fungal, plants and mammalian systems [6].

### The Arabidopsis thaliana

AMT1.1 is the first cloned and functionally characterized plant ammonium transporter through yeast complementation using mutant strain 26972c (*mep1 mep2 ura3*) [7]. cDNA library screening in this NH_4_^+^ uptake-deficient mutant identified the plant NH_4_^+^ transporter AtAMT1.1. The *Arabidopsis thaliana* genome encodes six AMT isoforms [8], while *Oryza sativa* possesses at least ten members, the latter could be classified into four phylogenetically distinct subtypes OsAMT1-4 [2,9]. As rice is primarily cultivated in anaerobic paddy soils where NH_4_^+^ predominates over NO_3_^-^ as the nitrogen source [10], this crop exhibits preferential ammonium uptake. Consequently, NH₄^+^ assimilation mechanisms have become a major research focus in rice physiology [11]. In rice, OsAMT1;1, OsAMT1;2, and OsAMT1;3 form a high-affinity ammonium transporter system [12,13]. These OsAMT1 members are polarly localized at root exodermis and collectively mediate the ammonium uptake under low ammonium concentration, and triple-gene knockout reduces root ammonium absorption by 95% [14]. Besides, *OsAMT1;1* is expressed in both roots and shoots, and induced by NH4^+^ but repressed under nitrogen starvation [12]. The overexpression of OsAMT1;1 enhances NH_4_^+^ permeability, promoting growth and yield under suitable NH_4_^+^ conditions [15]. Biochemically, OsAMT1;1 functions as a pH-independent ammonium uniporter with *Km* ≈ 110 μM for NH_4_^+^, subject to feedback inhibition by intracellular substrate accumulation [13]. Beyond plant systems, the crystal structures of AMT homologues from bacteria, archaea, yeast and human reveal conserved architecture in the AMT/MEP/Rh superfamily [16‒21]. These transporters form stable trimers where each monomer contains an extracellular vestibule that recruits NH_4_^+^ and function like a cation selective channel [22]. Substrate deprotonation at a conserved Twin-His motif enables NH_3_ transit through the central hydrophobic channel [16,22,23]. Cytosolic re-protonation occurs after passage through a second hydrophobic gate, driven by cytoplasmic pH gradients [24].

Ammonium transport is tightly regulated to prevent toxic accumulation. Intriguingly, distinct regulatory mechanisms have evolved across organisms. In bacteria, the trimeric PII-like protein GlnK directly blocks the AmtB channel during nitrogen sufficiency by binding its cytosolic pores [25,26]. In *Saccharomyces cerevisiae*, nitrogen limitation activates Npr1 kinase, which phosphorylates Mep2 at Ser457 to enhance transport activity [27]. Conversely, in higher plants, AMT proteins employ phosphorylation-based inhibition mechanisms at posttranscription level [28‒34]. For example, *Arabidopsis* CIPK23 kinase phosphorylates Thr460 of AtAMT1;1 to inhibit ammonium transport and modulate ammonium sensitivity [30,32]. Similarly, rice deploys OsACTPK1, a serine/threonine/tyrosine kinase, to phosphorylate OsAMT1;2 at residue Thr453 under sufficient ammonium supply [33]. The phosphorylation-mimetic substitution, by mutating conserved threonine residues to negative-charged aspartic acid, abolished ammonium transport activity in both yeast and *Xenopus* oocyte heterologous systems. Crucially, co-expression experiments reveal that phosphorylation of a single monomer is sufficient to inactivate the entire AMT1 trimer [28,29], indicating allosteric regulation through inter-subunit communication. Current models propose that homo-or hetero-trimerization enables physical interactions between the cytosolic C-terminal region of one monomer and adjacent monomer loops, facilitating conformational coupling and trans-activation dependent on phosphorylation [28,31], providing an explanation for rapid shutdown of NH_4_^+^ uptake to prevent toxicity. However, direct structural evidence for phosphorylation-induced inhibition mechanism remains elusive, and whether monomer phosphorylation induces autoinhibition concurrently requires further investigation.

In this study, we determined the cryo-electron microscopy (Cryo-EM) structures of OsAMT1;1 and its phosphomimetic variant OsAMT1;1-T452D. OsAMT1;1 forms stable trimer in lipid environment, with key functional motifs involved in NH_4_^+^ conduction demonstrating remarkable conservation across the AMT/MEP/Rh superfamily. Structural analysis combined with functional assays reveals that the T452D phosphomimetic stabilizes the closed state of the channel, supporting a mechanism for phosphorylation-induced inactivation within plant AMT1 subfamily.

## Materials and Methods

### Expression and purification of protein

The gene encoding *OsAMT1;1* was amplified from *Oryza sativa* cDNA. The fragments of *OsAMT1;1* with removal of the N-terminus (residues 2-19) was subcloned into modified pPICZ C vector (Invitrogen, Carlsbad, USA) within a TEV protease cleavage site and a C-terminal GFP-His tag. The T452D mutant was generated using two-step PCR. The plasmids were linearized with *Pme*I restriction enzyme (NEB, Ipswich, USA) and transformed into *Pichia pastoris* GS115 strain by electroporation (Bio-Rad, Hercules, USA). The positive clones were selected on agar plates containing 100 μg/mL zeocin (Invitrogen). For screening clones with high-expression level, about 20 yeast colonies were selected and grown overnight in minimal glycerol yeast nitrogen base (MGY) medium (10 mL), and then induced by adding 0.5% methanol in minimal methanol (MMH) medium for 48 h. Cell cultures were checked through the GFP fluorescence by confocal microscope. The clone with strongest fluorescence was selected and used for further large-scale expression.

To purify the proteins, 2.4L cultures were harvested and re-suspended in buffer A (20 mM Tris-HCl, pH 8.0,100 mM NaCl, and 5% glycerol) supplemented with 1 mM PMSF, 2 mM Mg^2+^ and 5 μg/mL DNase, and homogenized with UH-06 (Union-Biotech, Shanghai, China) for six times at 17,000 p.s.i. The lysate was centrifuged at 20,000 *g* for 20 min, and the supernatant was collected and ultracentrifuged at 38,000 r.p.m. (Beckman type 70 Ti rotor; Beckman Coulter, Brea, USA) for 1 h. The membrane fraction was re-suspended in buffer A and incubated with 1% (w/v) n-Dodecyl-β-D-Maltopyranoside (DDM; Anatrace, Maumee, USA) for 2 h at 4°C. Insoluble membranes were removed by centrifugation at 20,000 *g* for 1 h and the supernatant was added to NHS-activated Sepharose beads (GE Healthcare, Chicago, USA) coupled to GFP nanobody [35]. The beads were washed with buffer B (20 mM Tris-HCl, pH 8.0, 100 mM NaCl, 0.06% DDM, 0.006% cholesteryl hemisuccinate, and 2 mM DTT) and then incubated with TEV protease at 4°C overnight to remove C-terminal GFP tag. The protein was collected and concentrated for nanodisc reconstruction.

### Nanodisc reconstitution

Protein was reconstituted into lipid nanodisc following the published protocol [36]. Briefly, membrane scaffold protein MSP2N2 was expressed in *E. coli* BL21(DE3) and purified. 1-palmitoyl-2-oleoyl-sn-glycero-3-phosphoglycerol (POPG; Avanti, Alabaster, USA) solubilized in chloroform was dried under nitrogen gas and dissolved in sodium cholate buffer at a concentration of 10 mM. Protein, MSP2N2 and POPG were mixed at a molar ratio of 1:3:125 and incubated at 4°C for 1 h. Detergents were removed by additional Bio-Beads SM2 (Bio-Rad) to a concentration of 0.5 g/mL at 4°C overnight. The supernatant was collected and further purified by size-exclusion chromatography (Superdex 200 10/300 GL column; GE Healthcare) with buffer C (20 mM Tris-HCl, pH 8.0, and 100 mM NaCl). The peak fractions were concentrated to about 2 mg/mL for sample preparation.

### Cryo-EM sample preparation and data collection

To prepare samples for cryo-EM analysis, 3.5 μL of protein was applied to glow-discharged carbon grids (Quantifoil Au R1.2/1.3 300 mesh). Grids were blotted for 3 s with Grade 597 Filter Paper (Whatman, Kent, USA) and plunge-frozen in liquid ethane cooled by liquid nitrogen using a Vitrobot Mark IV (Thermo Fisher Scientific, Waltham, USA) at 8°C and 100% humidity. The grids were stored in liquid nitrogen until further use.

The data of OsAMT1;1 wild-type (WT) was collected on a Titan Krios (Thermo Scientific Scientific) transmission electron microscope operated at 300 kV and equipped with a K3 direct electron detector (Gatan, Pleasanton, USA). The movie stacks were automatically acquired in super-resolution mode (81,000×, magnification) using SerialEM software, with a set defocus range of -1.2 to -2.2 μm. Images were recorded for 2.5 s exposures in 32 subframes to give a total dose of 50 e^−^/Å^2^.

The data of OsAMT1;1 T452D variant was collected on a Titan Krios transmission electron microscopy operated at 300 kV and equipped with a K3 direct electron detector and GIF Quantum energy filter. The movie stacks were recorded in the super-resolution mode at nominal magnification of 81,000× with a calibrated pixel size of 0.539 Å. The defocus range was from -1.3 to -1.8 μm. Each stack of 32 frames was exposed for 2.56 s with a total dose of 50 e^−^/Å^2^. AutoEMation was used for automated data collection [37].

### Data processing

The data processing of OsAMT1;1 WT and T452D was performed using RELION [38] and the workflow and statistics are summarized in **Supplementary Figures S1‒2** and **Supplementary Table S1**. Raw movie stacks were first motion-corrected using MotionCorr2 [39] and the CTF parameters were estimated with Gctf [40].

After automatically pick selection, particles were extracted for each dataset. Junk particles were removed by multiple rounds of 2D classification. The initial 3D model was generated in RELION. Subsequent 3D classification was performed to clean the particles, and the best representative class was selected for 3D Auto-refinement. For the data of OsAMT1;1 WT, the application of C3 symmetry and post-processing results in a final map at the resolution of 3.82 Å. For the data of OsAMT1;1 T452D, the best class generate a high-quality map at the resolution of 2.44 Å after contrast transfer function (CTF) refinement with C3 symmetry applied. Post-processing was performed to further process particles, reaching a resolution of 2.19 Å. The overall resolutions were estimated based on the gold-standard Fourier shell correlation = 0.143 criterion. Local resolution estimation was calculated using RELION.

### Model building and structure refinement

The atomic models of OsAMT1;1 WT and T452D were built using the predicted model by trRosetta [41] as a template. The predicted model was docked into the density map and manually adjusted and rebuilt by COOT [42]. The resulting model was refined using real_space_refine module of PHENIX [43] with secondary structure and geometry restraints. The structures were validated with MolProbity [44], and refinement statistics can be found in **Supplementary Table S1**. All structure figures were prepared in PyMOL (Schrödinger) and ChimeraX [45].

### Yeast growth complementation assays

The fragments of OsAMT1;1 WT and mutants were cloned into yeast expression vector pDR196. The plasmid was transformed into yeast strain 31019b (*Δmep1 mep2::LEU2 mep3::KanMX2 ura3*) [46], and transformants were selected on YNB plates containing 0.17 % Yeast Nitrogen Base (YNB) without amino acids and ammonium sulfate (Difco) and supplemented with 2% (w/v) glucose and 1 mM arginine. Positive clones were grown in liquid arginine medium at 30°C for 2 days. Cells were collected, washed and re-suspended in sterile water, diluted 10^1^-10^4^ and dropped on solid YNB medium supplemented with 1 mM arginine or 1mM ammonium chloride as the sole N sources, buffered with 20 mM MES/Tris, pH 6.0. It should be noted that wild type and mutants were cloned into vector pDR196 with C-terminal GFP for fluorescence microscope observation. All experiments included three independent biological replicates.

## Results

### Phosphoregulatory conservation in OsAMT1;1

Sequence analysis revealed that the phosphorylation sites in rice and *Arabidopsis* AMTs are conserved among plant species (**Figure 1A**). To explore the functional conservation, we generated two phospho-state variants of OsAMT1;1, T452A mimicking the de-phosphorylated state, and T452D mimicking the constitutive phosphorylated state. In the ammonium-uptake-deficient yeast strain 31019b (*mep1Δ mep2Δ mep3Δ ura3*) [46], T452A mutant restored growth similar to wild-type OsAMT1;1 on culture medium containing ammonium as the sole N source, whereas T452D mutant showed growth inhibition similar to empty vector control (**Figure 1B**). Fluorescence of GFP-tagged OsAMT1;1 variants confirmed the unaffected stability and subcellular localization (**Supplementary Figure S3A**). The substitution of Thr452 to aspartic acid lost complementation capacity, indicating that the conserved phosphorylation inactivates OsAMT1;1, extending this regulatory mechanism to more family members [29,32,33,47].

**Figure 1.**
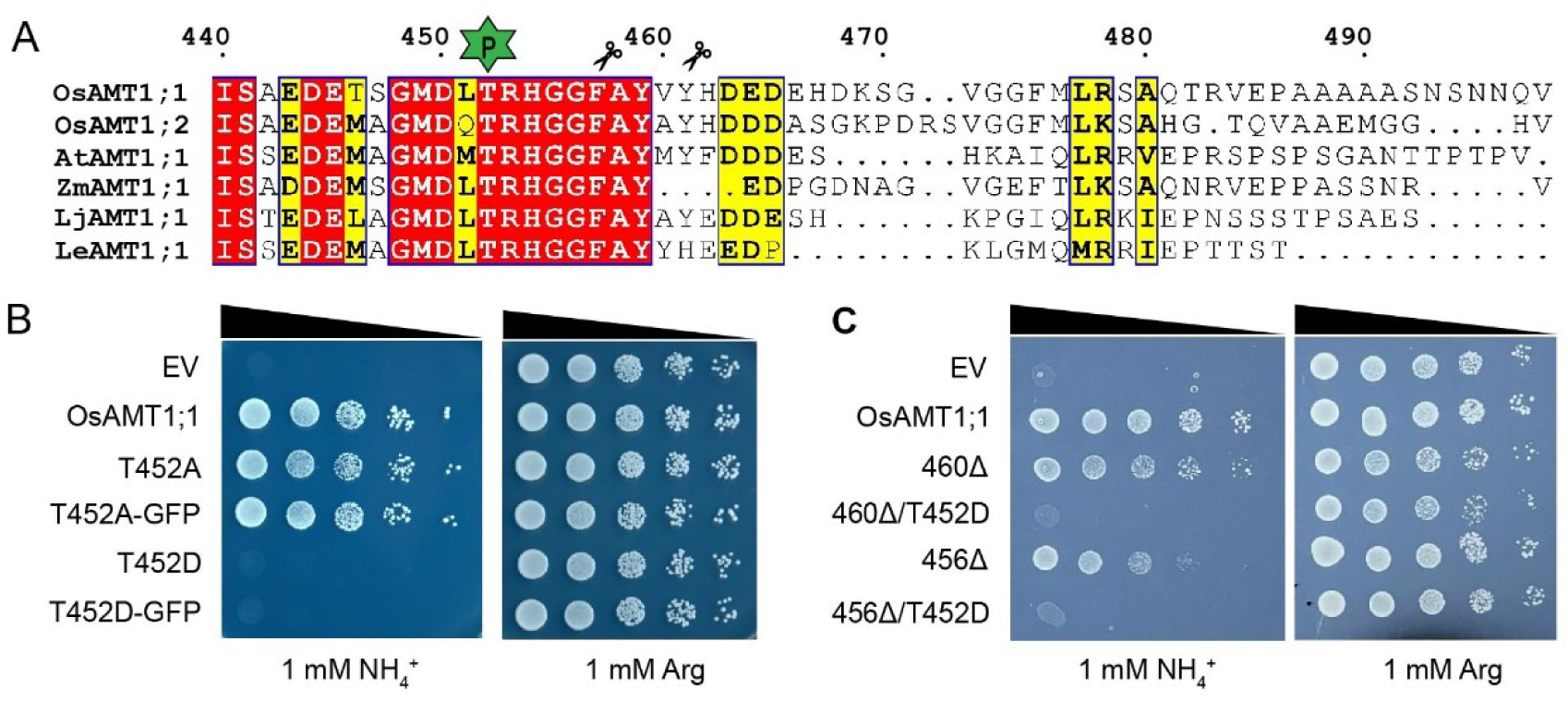
Functional characterization of OsAMT1;1 variants. (A) Multiple sequence alignment of C-terminal region from OsAMT1;1 and AMT proteins. Invariant residues are highlighted with red box; conserved residues are highlighted by yellow box. The phosphorylation site and C-terminal truncation sites (460Δ/456Δ, deleting residues after Val460/Gly456) are indicated. *Oryza sativa* AMT1;1, Uniprot Q7XQ12; *Oryza sativa* AMT1;2, Uniprot Q6K9G1; *Arabidopsis thaliana* AMT1;1, Uniprot P54144; *Zea mays* AMT1;1, Uniprot B4FV06; *Lotus japonicus* AMT1;1, Uniprot Q9FSH3; *Lycopersicon esculentum* AMT1;1, Uniprot P58905. (B,C) Complementation by pDR196-expressed OsAMT1;1 phosphorylation site mutants (B) and C-terminal region truncation mutants alone or in combination with T452D (C) supplemented with 1 mM arginine or 1 mM ammonium chloride. EV, empty vector.

To investigate the potential role of the flexible cytosolic C terminal region in phosphorylation-related regulation, we generated two C-terminal truncations of the full-length *OsAMT1;1* to eliminate the possible interaction between C-terminus and the cytosolic loops from the neighboring monomer. Yeast complementation assays showed that both the 460Δ (deleting residues after Val460) and 456Δ truncations remained functional, indicating that the C-terminal region is dispensable for ammonium transport (**Figure 1A, C**). However, introducing the phosphomimetic mutation T452D into these truncated variants abolished their complementation capacity, which is consistent with the inactivation observed in the full-length T452D variant (**Figure 1C**). These results provide the evidence that phosphorylation-mediated inactivation can occur in the absence of cytosolic C-terminal region.

### Overall structure of OsAMT1;1

To elucidate the phosphorylation-mediated inhibition mechanism, we purified the protein of OsAMT1;1 in *Pichia pastoris* for structural analysis, and the N-terminal 19 residues were truncated to facilitate protein expression. Functional complementation assays confirmed that this truncation retained the ability to rescue the growth defect in yeast strain 31019b (**Supplementary Figure S3B**). Both the wild-type and the phosphomimetic T452D variant were purified in detergent and reconstituted into lipidic nanodiscs for cryo-EM analysis (as detailed in the **Materials and Methods**). We determined the structure of the wild-type OsAMT1;1 at an overall resolution of 3.8 Å (**Supplementary Figure S1**), and the T452D variant at 2.2 Å resolution, both under C3 symmetry (**Supplementary Figure S2**). The electron density maps are well-defined and allow us for accurate model building of the majority of the protein (**Supplementary Figure S4**), except for the deleted N-terminus and the disordered C-terminal region (from Val460 to the end).

OsAMT1;1 forms a stable homotrimer in a lipidic environment (**Figure 2A**). Each monomer consists of 11 transmembrane helices surrounding a central channel for ammonium transport (**Figure 2A, B**). Key structural features of OsAMT1;1 are highly conserved compared to other known structures of ammonium transporter [16‒18,20,21]: specifically, residues Ser225 and Trp168 form an extracellular ammonium recruitment site; residues Phe127 and Phe251 form a hydrophobic gate that separates the periplasmic vestibule from the hydrophobic lumen; and the twin-His motif, formed by residues His199 and His366, locates at the center of the narrow hydrophobic channel, and is responsible for substrate selection, NH_4_^+^ deprotonation and NH_3_ conduction (**Figure 2C**). At the cytosolic exit of the channel, a second constriction consists of residues Leu56, Asn312 and Leu360 separates the hydrophobic channel from the intracellular vestibule (**Figure 2C**). The strong structural conservation of these functional motifs suggests that the core transport mechanism of the AMT/MEP/Rh superfamily is preserved in OsAMT1;1.

**Figure 2.**
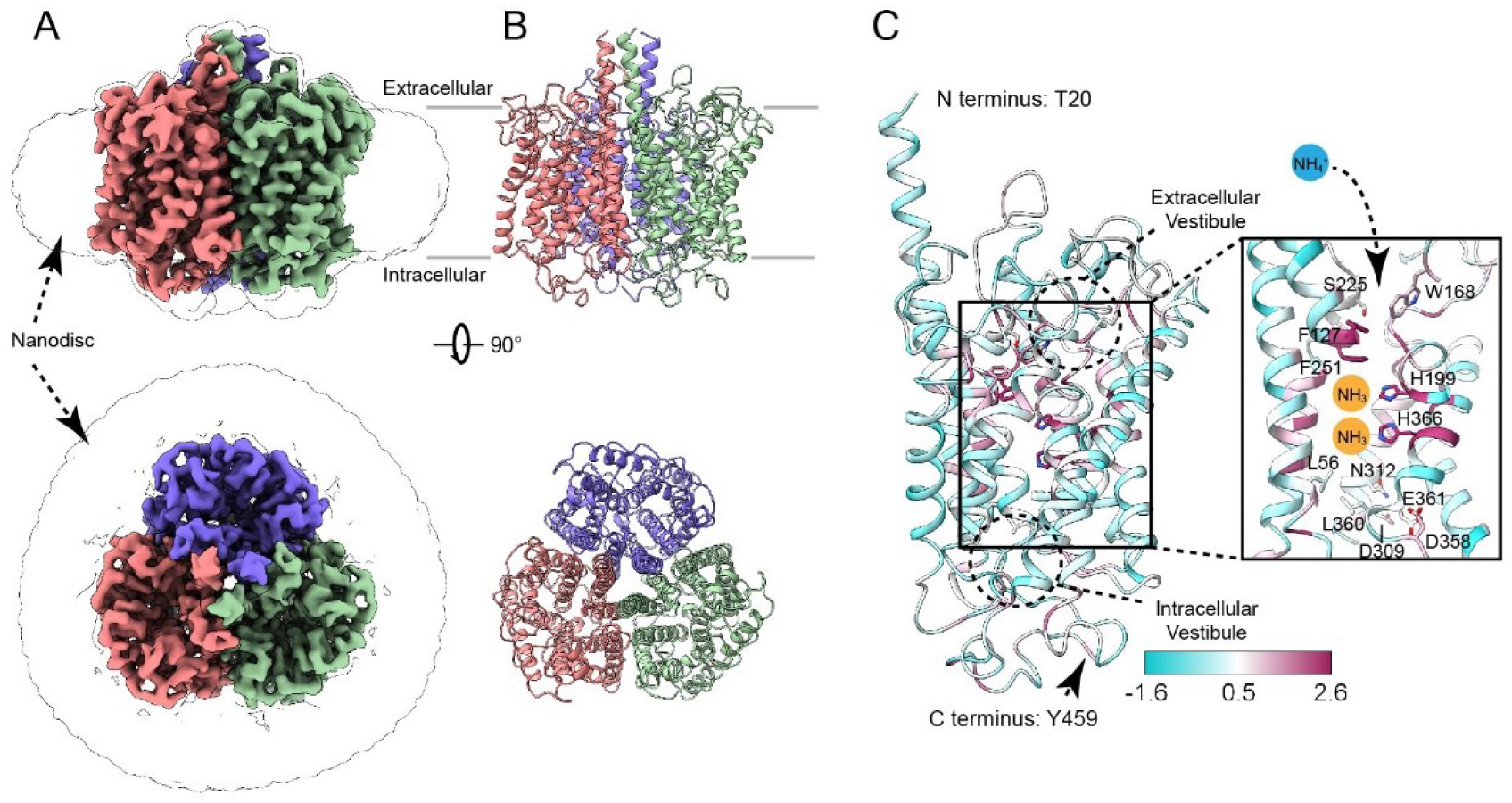
Overall structure of OsAMT1;1. (A,B) Cryo-EM density map (A) and overall structure (B) of OsAMT1;1. The three monomers of OsAMT1;1 are colored in pink, green and purple, respectively. (C) Key functional features of OsAMT1;1. The OsAMT1;1 monomer is shown in cartoon and colored by sequence conservation as calculated by ConSurf ranging from cyan to maroon. Inset, a zoom-in view with functional motifs shown as sticks and ammonium conduction pathway indicated.

The overall architecture of OsAMT1;1 is similar to AmtB from *E. coli*, with a Cα r.m.s.d. of 4.4 Å over 377 aligned residues. While the transmembrane domain closely resembles that of AmtB, obvious structural differences are observed in the extracellular and intracellular loop regions (**Figure 3A**). Notably, the N-terminus of TM1 is elongated compared to AmtB, which expands the interface between monomers to stabilize the trimer, supported by a previous study on plant AMT proteins [48]. Electrostatic surface analysis reveals two negatively charged cavities located on opposite sides of OsAMT1;1 (**Figure 3B**), suggesting a pathway for substrate conduction. The extracellular cavity is more contractive and distinctly shaped than that of AmtB (**Figure 3C**). On the intracellular side, the cavity connects directly to the channel exit (**Figure 3B**) and is mainly formed by the charged residues Asp309, Asp358 and Glu361 (**Figure 2C**), likely constituting a pocket for NH_4_^+^ release.

**Figure 3.**
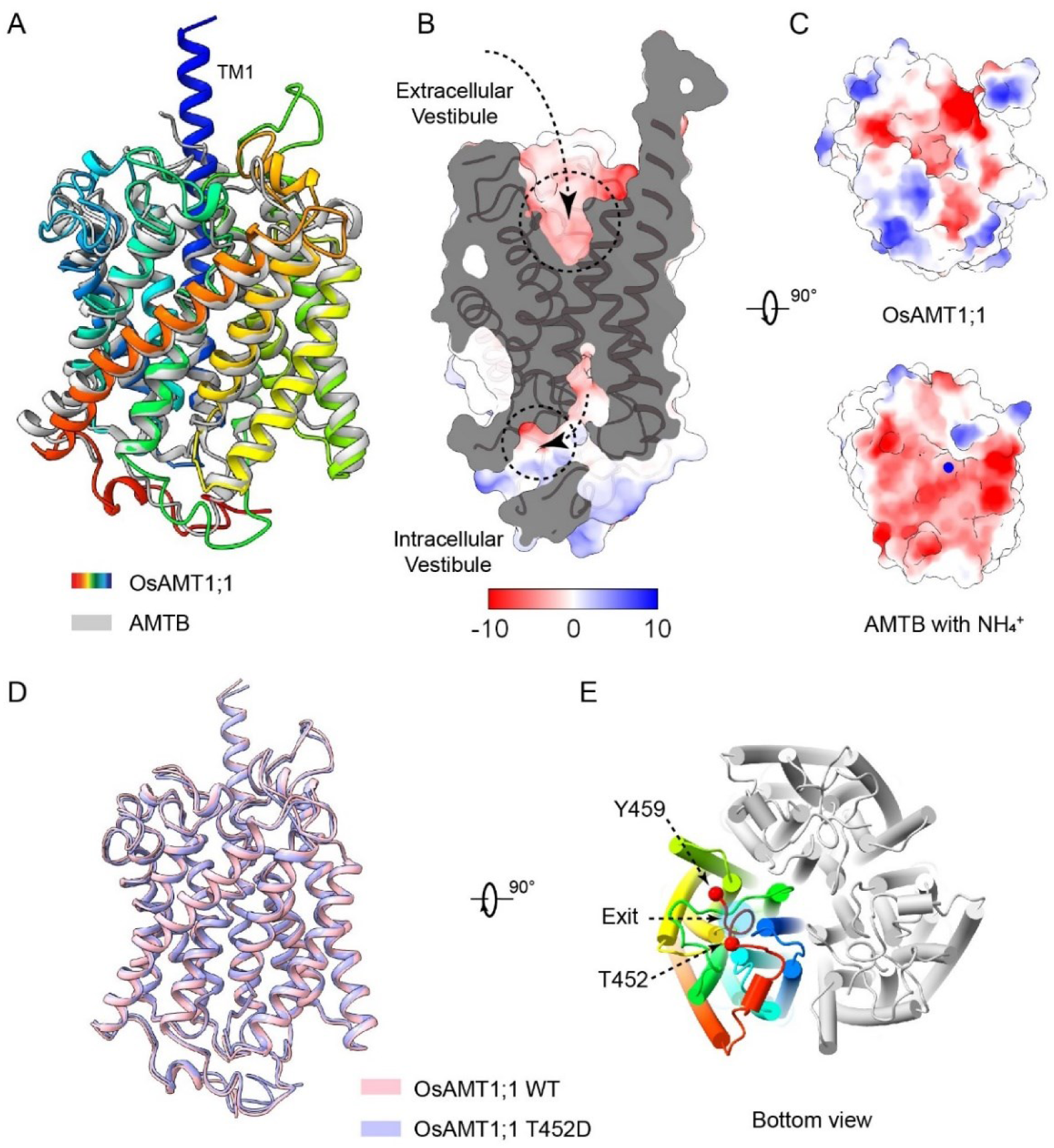
Structural comparison of OsAMT1;1 WT and T452D. (A) Structural superposition of OsAMT1;1 and AMTB with NH_4_^+^ (PDB 1U7G). OsAMT1;1 and AMTB structures are colored in rainbow and grey. (B) Electrostatic surface analysis of OsAMT1;1 monomer. The side view of the cut-open electrostatic surface of OsAMT1;1 showing two negatively charged cavities, with the potential substrate pathway indicated by arrows. The transmembrane helices are shown as ribbons. (C) Extracellular vestibule of OsAMT1;1 in comparison with AMTB. (D) Structural superposition of OsAMT1;1 WT and T452D monomer. OsAMT1;1 WT and T452D structures are colored in pink and purple, respectively. (E) Intracellular view of the OsAMT1;1 T452D trimeric structure. One of the monomers is colored in rainbow and the others in grey. Residues T452D and Y459 are shown as spheres.

To elucidate the structural mechanism underlying phosphorylation-mediated regulation in the plant AMT1 subfamily, we compared the cryo-EM structures of wild-type OsAMT1;1 and the phosphomimetic T452D variant. Overall, the two structures are very similar, with a Cα r.m.s.d. of 0.69 Å over 435 aligned residues (**Figure 3D**). In both cases, the cytosolic C-terminal tail beyond Tyr459 is disordered and could not be modeled. No clear interaction was observed between the resolved cytosolic C-terminus (up to Tyr459) and the cytosolic loops of neighboring monomers (**Figure 3E**). The phosphorylation site Thr452 is situated near the beginning of the partially disordered C-terminal tail, distal to the channel exit but in proximity to intracellular loop regions (**Figure 3E**). Consistent with our functional data showing that truncated variants (460Δ/T452D and 456Δ/T452D) are still inactivated by the T452D mutation, these structural analyses suggest that the phosphorylation regulatory mechanism involves more functional elements in addition to the cytosolic C-terminal region.

### Channel state of OsAMT1;1 wild-type and T452D variant

The T452D variant of OsAMT1;1 represents an inactivated channel state, while the high structural similarity between the wild-type and T452D structures suggests that the wild-type protein also adopts a closed state under the conditions examined. This is consistent with the fact that the proteins were expressed in a rich ammonium medium, likely favoring a closed state. We next superimposed the structures of OsAMT1;1-T452D, and AmtB in apo state and inhibitory state (in complex with the inhibitory protein GlnK) (**Figure 4A**). Although the transmembrane domains showed no major conformational differences, the channel of AmtB in the AmtB-GlnK complex is directly blocked by GlnK, whose extended T-loop inserts Arg47 deeply into the cytoplasmic vestibule of AmtB channel (**Figure 4B**). This vestibule, formed primarily by TM8, TM9 and TM10, undergoes a conformational rearrangement in intracellular loop 5 (ICL5, located between TM9 and TM10) to accommodate the side chain of Arg47 when compared to AmtB in apo state. Interestingly, in both OsAMT1;1 wild-type and T452D structures, the conformation of ICL5 closely resembles that in the AmtB-GlnK blocked state (**Figure 4B, C**), further supporting the conclusion that both structures are in a closed sate.

**Figure 4.**
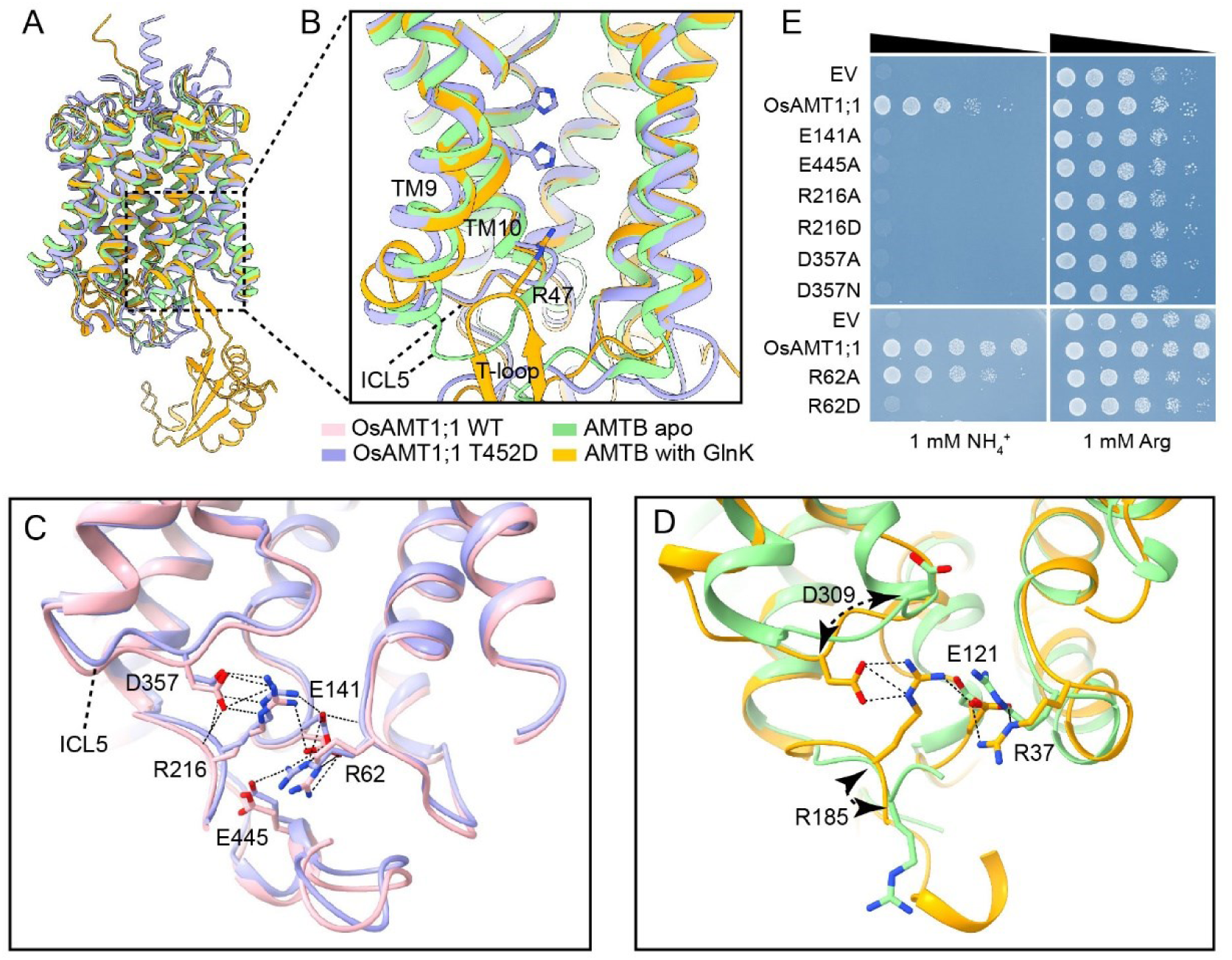
OsAMT1;1 channel state and salt-bridge network. (A) Structural superposition of OsAMT1;1 T452D, AMTB apo (PDB 1U77) and AMTB-GlnK (PDB 2NUU) monomers. OsAMT1;1 T452D, AMTB apo and AMTB-GlnK structures are colored in purple, green and yellow, respectively. (B) A zoom-in view of the intracellular vestibule in (A). The residue Arg47 from AMTB-GlnK is shown as sticks. TM9 and TM10 are labelled, while TM8 is omitted in this view. (C) Intracellular vestibule of OsAMT1;1 WT and T452D structures. The salt-bridge network is indicated and the coordinating residues are shown as sticks. (D) Intracellular vestibule of AMTB apo and AMTB-GlnK. (E) Complementation by pDR196-expressed OsAMT1;1 mutants supplemented with 1 mM arginine or 1 mM ammonium chloride. EV, empty vector.

In yeast, Mep2 from *Saccharomyces cerevisiae* and *Candida albicans* also adopt a closed, inactive conformation under nitrogen-sufficient conditions [21]. Unlike their plant and bacteria orthologues, the longer ICL1 (located between TM1 and TM2) and the cytoplasmic end of TM2 in CaMep2 shows inward movement, and the residue Tyr49 protruded from TM2 forms hydrogen bonds with the twin-His motif to block the channel (**Supplementary Figure S5A**). Therefore, the regulation mechanism of ammonium transport in yeast is distinct from that observed in plants.

It is noteworthy that structural analysis revealed an electrostatic contact network within the intracellular vestibule of OsAMT1;1. In both wild-type and T452D structures, Asp357 from ICL5 forms a salt bridge with Arg216, which in turn interacts with Glu141. Glu141 further engages in a salt bridge with Arg62, and Arg62 also forms a salt bridge with Glu445 (**Figure 4C**). A similar interaction pattern is observed in the AmtB-GlnK complex, involving the corresponding residues Asp309, Arg185, Glu121, and Arg37. In contrast, these electrostatic interactions are absent in the apo state of AmtB, as the conformational change of intracellular loops disrupts the salt bridge network, especially the reorientation of Asp309 and Arg185 (**Figure 4D**). This suggests that the formation and dissolution of these salt bridges are associated with channel gating. Functional assays confirmed that mutation of residues within this interaction network could abolish the complementation capacity of OsAMT1;1 (**Figure 4E**). The sequence conservation analysis shows that all these residues are highly conserved across the AMT/MEP/Rh superfamily (**Supplementary Figure S5B**), underscoring their essential role in ammonium transport function.

### Phosphorylation stabilizes channel in closed state

Notably, Arg62 is located in proximity to Thr452, though it is not a core component of the electrostatic network. While the R62A mutation retained the ability to restore yeast growth, substitution to aspartate (R62D) yielded a non-functional transporter (**Figure 4E**), mirroring the behavior of the T452A and T452D mutants. Although the phosphorylation site Thr452 lies near a disordered region, and its electron density lacked high precision (while detectable in both OsAMT1;1 WT and T452D maps), the density of ICL5 and residues within the salt bridge network was well resolved (**Supplementary Figure S5C, D**). The structural comparison suggests that the salt-bridge network is better resolved and likely stabilized in the T452D structure (**Figure 5A**). The upward movement of the Arg62 in the T452D structure is likely to accommodate the negative charge introduced by the phosphorylation. This shift appears to propagate throughout the interaction network, as all salt-bridging residues exhibited a similar upward movement in the T452D structure, as well as the ICL5 loop (**Figure 5A**). The enhanced stability of the salt bridge interactions in the T452D mutant is further supported by the higher resolution achieved in its structure compared to the wild-type. These findings suggest that the phosphomimetic T452D variant stabilizes the closed state of the channel by reinforcing the salt bridge network within the intracellular vestibule.

**Figure 5.**
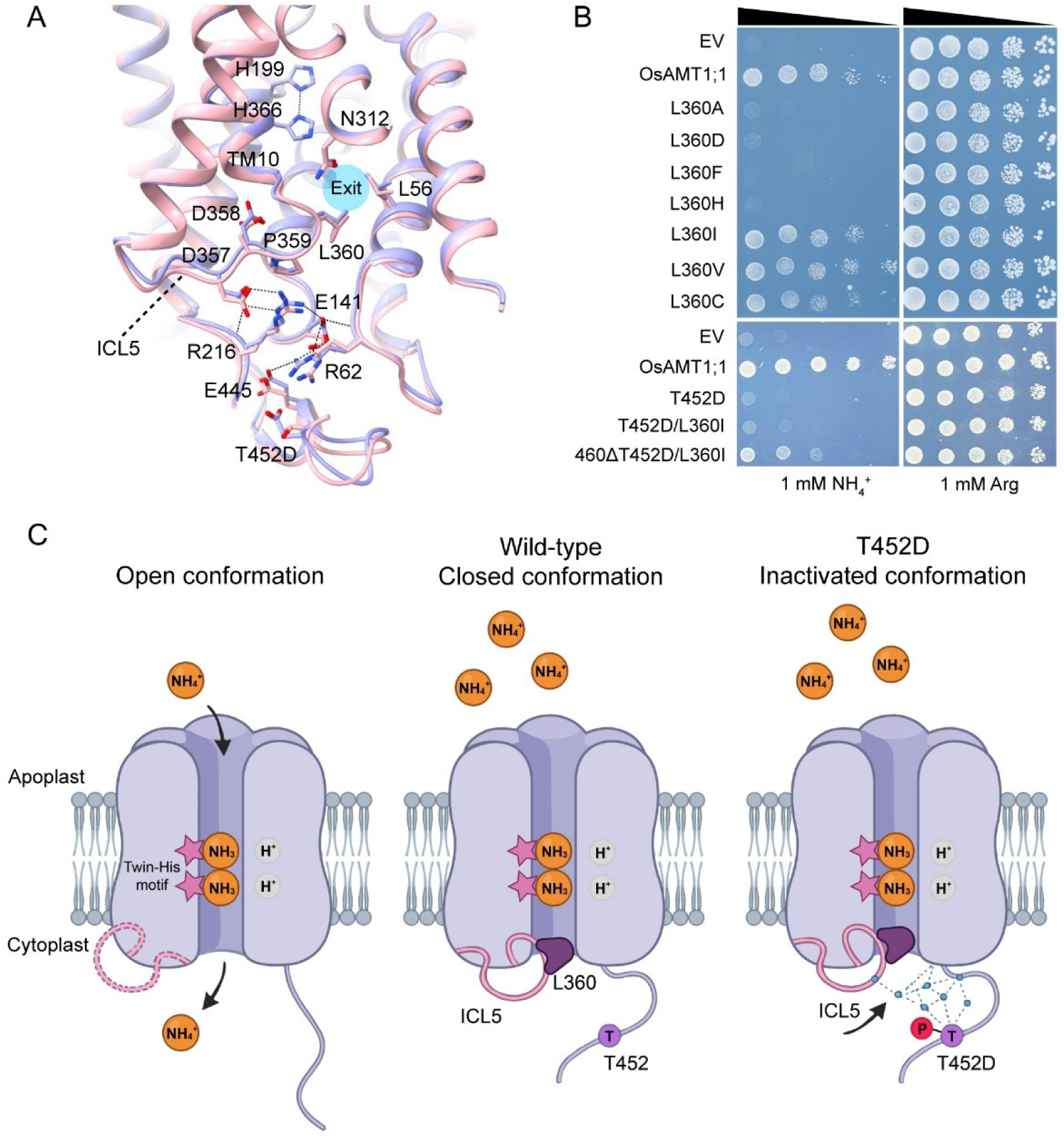
Proposed regulatory mechanism of OsAMT1;1. (A) Intracellular vestibule of OsAMT1;1 WT and T452D structures. The salt-bridge network is indicated and the coordinating residues are shown as sticks. (B) Complementation by pDR196-expressed OsAMT1;1 mutants alone or in combination with T452D supplemented with 1 mM arginine or 1 mM ammonium chloride. EV, empty vector. (C) Proposed regulatory mechanism of OsAMT1;1. The OsAMT1;1 structure and key features are condensed for illustrative purposes. Only one subunit of the functional transporter is shown for clarity. In this model, the substrate pathway of OsAMT1;1 is blocked by plug Leu360 (middle, closed conformation). The phosphorylation at the conserved residue Thr452 stabilizes an intricate salt-bridge network within the intracellular vestibule of the transporter. This network acts collectively to induce an upward conformational shift in ICL5, a cytoplasmic loop that functions as a gating loop. The movement of ICL5 is subsequently transmitted to the hydrophobic plug near the cytoplasmic exit, thereby occluding the channel exit and abolishing ammonium transport (right, inactivated conformation). In the absence of phosphorylation and under low ammonium conditions, the salt-bridge network is dynamic or disordered, allowing ICL5 to adopt a relaxed conformation, opening the cytoplasmic exit and permitting ammonium release (left, hypothetical open conformation).

The movement of ICL5 does not induce the conformational change of TM10. Instead, the linker region between ICL5 and TM10 protrudes into the channel exit like a plug. Leu360 serves as the central residue of this plug and, together with Leu56 and Asn312, constitutes a hydrophobic barrier that separates the main channel from the intracellular vestibule (**Figure 5A**). Functional assays revealed that the mutation of Leu360 to Alanine, Aspartic acid, Histidine or the aromatic residue Phenylalanine disrupted transport activity, whereas substitution with similar residues such as Isoleucine, Valine or Cysteine did not impair function (**Figure 5B**). Given that Leucine is among the strongest residues forming α-helical and β-strand, these results indicate the critical role of Leu360 in maintaining the hydrophobicity and structural integrity of the channel exit, thereby stabilizing the plug architecture.

In conclusion, based on structural and functional analyses, we propose a regulatory mechanism for phosphorylation-induced inactivation of OsAMT1;1. Phosphorylation of the conserved threonine stabilizes a network of salt bridges within the intracellular vestibule. ICL5 acts as a gating loop that transmits the phosphorylation-induced conformational change to Leu360, maintaining the channel in a stably closed state. Furthermore, the high conservation of Leu360 and the network-forming residues (**Supplementary Figure S5B**), relative to Thr452, suggests that phosphorylation-induced inactivation in plants may have evolved more recently than the ancestral transport mechanism shared within the AMT/MEP/Rh superfamily.

### L360I partially rescues phosphomimetic variant upon C-terminal truncation

To further investigate whether Leu360 functions as the key effector in phosphorylation-induced channel inactivation, we attempted to restore transport activity in the T452D variant by substituting Leu360 with smaller side-chain residues. Given that the L360A mutation alone abolishes protein function, we generated the double mutants T452D/L360I, T452D/L360V and T452D/L360C. Yeast complementation assays showed that none of these variants recovered ammonium transport (**Supplementary Figure S3C**), despite exhibiting expression and subcellular localization comparable to the wild-type protein.

We have identified a regulatory mechanism for phosphorylation-mediated inactivation that operates within AMT1 monomer. This finding does not preclude the existence of additional regulation mediated by the C-terminal region across monomers. Instead, we propose that these two regulatory mechanisms may function synergistically to enable rapid and efficient prevention of excess ammonium uptake in plants. This model could explain the inability of the L360I, L360V or L360C substitutions to restore function in the T452D background. To test the hypothesis, we removed the C-terminal region and constructed three additional double mutants: 460Δ/T452D/L360I, 460Δ/T452D/L360V and 460Δ/T452D/L360C. Intriguingly, the triple mutant 460Δ/T452D/L360I partially rescued yeast growth, although its transport activity remained lower than that of the wild type (**Figure 5B**).

The partial function rescue of the T452D variant by L360I upon removal of the C-terminal disordered tail confirms the critical role of L360 in phosphorylation-induced channel blockage. These results are consistent with a working hypothesis in which phosphorylation regulates AMT activity through two cooperating mechanisms: intra-monomer salt bridge stabilization and inter-monomer C-terminal interactions, enabling rapid inhibition of AMTs.

## Discussion

The phosphorylation-mediated regulation of ammonium influx by AMTs is essential for plant growth and stress responses. While the function of the AMTs has been established, the structural and mechanistic basis of its phosphorylation induced inhibition remains elusive. Our cryo-EM structures of the rice ammonium transporter OsAMT1;1 and its phosphomimetic variant T452D, combined with functional analyses, suggest a regulatory mechanism for the phosphorylation-induced inactivation of plant AMT1 subfamily. In this model, phosphorylation at the conserved residue Thr452 stabilizes an intricate salt-bridge network within the intracellular vestibule of the transporter. This network includes key residues such as Arg62, Glu141, Arg216, and Asp357, which act collectively to induce an upward conformational shift in ICL5, a cytoplasmic loop that functions as a gating loop. The movement of ICL5 is subsequently transmitted to the hydrophobic plug residue Leu360 near the cytoplasmic exit, thereby occluding the channel exit and abolishing ammonium transport (**Figure 5A, C**). Our data highlight the intra-monomer gating by the ICL5 loop for transporter inactivation, while not excluding the role of C-terminal region in fine-tuning the response or in mediating additional forms of regulation [28,31]. This mechanism may cooperate with inter-monomeric regulatory mechanisms involving the C-terminal region to enable rapid inhibition of ammonium uptake, and the precise role of the C-terminal domain requires further investigation.

Notably, the conformation observed in both the wild-type and T452D mutant structures corresponds to a closed state, likely reflecting the ammonium-replete conditions under which the proteins were expressed. A critical unanswered question is how the open state of OsAMT1;1 is structurally realized. We hypothesize that in the absence of phosphorylation and under low ammonium conditions, the salt-bridge network is dynamic or disordered, allowing ICL5 to adopt a relaxed conformation. This would withdraw Leu360 from the pore, opening the cytoplasmic exit and permitting ammonium release (**Figure 5C**, left). The transition between open and closed states may involve the disruption of specific salt bridges and a slight inward flexing of TM helices adjacent to the conduction path.

In conclusion, our work provides a structural and mechanistic basis for understanding phosphorylation-mediated inactivation of plant AMT1 transporters, in contrast to the phosphorylation activation of fungal homologs. It reveals a regulatory mechanism that allows plants to rapidly inhibit ammonium uptake in response to sufficient nitrogen availability to prevent toxicity.

## Data Availability

The cryo-EM density maps of OsAMT1;1 WT and T452D have been deposited in the Electron Microscopy Data Bank (EMDB) under accession codes EMD-67958 and EMD-67959. The corresponding atomic coordinates have been deposited in the Protein Data Bank (PDB) under accession codes PDB 21SC and 21SD.

## Acknowledgements

We thank the staff members at the cryo-EM centers of Westlake University and Peking University for their technical assistance on cryo-EM data collection. We thank Dr Lixing Yuan (China Agricultural University) for providing the yeast strain. We thank Dr Xinqi Gong (Renmin University of China), Dr Jian Feng Ma and Dr Noriyuki Konishi (Okayama University) for helpful discussions.

## Author Contributions

FY, XH, and XZ designed the experiments. XH, FY, SW and NA carried out protein expression, sample preparation, cryo-EM data collection and yeast growth assays. PZ and JH contributed to data analysis. FY, XH and XZ wrote the manuscript. XZ and JH conceived the project. All authors edited and approved the manuscript.

## References

1. Xu, G., X. Fan, and A.J. Miller, Plant nitrogen assimilation and use efficiency. Annu Rev Plant Biol, 2012. 63: p. 153–82.

2. Xuan, W., T. Beeckman, and G. Xu, Plant nitrogen nutrition: sensing and signaling. Curr Opin Plant Biol, 2017. 39: p. 57–65.

3. Bloom, A.J., S.S. Sukrapanna, and R.L. Warner, Root respiration associated with ammonium and nitrate absorption and assimilation by barley. Plant Physiol, 1992. 99(4): p. 1294–301.

4. von Wiren, N., et al., The molecular physiology of ammonium uptake and retrieval. Curr Opin Plant Biol, 2000. 3(3): p. 254–61.

5. Hao, D.L., et al., Function and Regulation of Ammonium Transporters in Plants. Int J Mol Sci, 2020. 21(10).

6. Loque, D. and N. von Wiren, Regulatory levels for the transport of ammonium in plant roots. J Exp Bot, 2004. 55(401): p. 1293–305.

7. Ninnemann, O., J.C. Jauniaux, and W.B. Frommer, Identification of a high affinity NH4+ transporter from plants. EMBO J, 1994. 13(15): p. 3464–71.

8. Ludewig, U., B. Neuhauser, and M. Dynowski, Molecular mechanisms of ammonium transport and accumulation in plants. FEBS Lett, 2007. 581(12): p. 2301–8.

9. von Wittgenstein, N.J., et al., Evolutionary classification of ammonium, nitrate, and peptide transporters in land plants. BMC Evol Biol, 2014. 14: p. 11.

10. Li, Y.L., X.R. Fan, and Q.R. Shen, The relationship between rhizosphere nitrification and nitrogen-use efficiency in rice plants. Plant Cell Environ, 2008. 31(1): p. 73–85.

11. Wang, M.Y., et al., Ammonium Uptake by Rice Roots (I. Fluxes and Subcellular Distribution of 13NH4+). Plant Physiol, 1993. 103(4): p. 1249–1258.

12. Sonoda, Y., et al., Distinct expression and function of three ammonium transporter genes (OsAMT1;1-1;3) in rice. Plant Cell Physiol, 2003. 44(7): p. 726–34.

13. Yang, S., et al., The rice OsAMT1;1 is a proton-independent feedback regulated ammonium transporter. Plant Cell Rep, 2015. 34(2): p. 321–30.

14. Konishi, N. and J.F. Ma, Three polarly localized ammonium transporter 1 members are cooperatively responsible for ammonium uptake in rice under low ammonium condition. New Phytol, 2021. 232(4): p. 1778–1792.

15. Ranathunge, K., et al., AMT1;1 transgenic rice plants with enhanced NH4(+) permeability show superior growth and higher yield under optimal and suboptimal NH4(+) conditions. J Exp Bot, 2014. 65(4): p. 965–79.

16. Khademi, S., et al., Mechanism of ammonia transport by Amt/MEP/Rh: structure of AmtB at 1.35 A. Science, 2004. 305(5690): p. 1587–94.

17. Zheng, L., et al., The mechanism of ammonia transport based on the crystal structure of AmtB of Escherichia coli. Proc Natl Acad Sci U S A, 2004. 101(49): p. 17090–5.

18. Andrade, S.L., et al., Crystal structure of the archaeal ammonium transporter Amt-1 from Archaeoglobus fulgidus. Proc Natl Acad Sci U S A, 2005. 102(42): p. 14994–9.

19. Lupo, D., et al., The 1.3-A resolution structure of Nitrosomonas europaea Rh50 and mechanistic implications for NH3 transport by Rhesus family proteins. Proc Natl Acad Sci U S A, 2007. 104(49): p. 19303–8.

20. Gruswitz, F., et al., Function of human Rh based on structure of RhCG at 2.1 A. Proc Natl Acad Sci U S A, 2010. 107(21): p. 9638–43.

21. van den Berg, B., et al., Structural basis for Mep2 ammonium transceptor activation by phosphorylation. Nat Commun, 2016. 7: p. 11337.

22. Ariz, I., et al., Nitrogen isotope signature evidences ammonium deprotonation as a common transport mechanism for the AMT-Mep-Rh protein superfamily. Sci Adv, 2018. 4(9): p. eaar3599.

23. Wang, S., et al., Ammonium transporters achieve charge transfer by fragmenting their substrate. J Am Chem Soc, 2012. 134(25): p. 10419–27.

24. Williamson, G., et al., A two-lane mechanism for selective biological ammonium transport. Elife, 2020. 9.

25. Durand, A. and M. Merrick, In vitro analysis of the Escherichia coli AmtB-GlnK complex reveals a stoichiometric interaction and sensitivity to ATP and 2-oxoglutarate. J Biol Chem, 2006. 281(40): p. 29558–67.

26. Gruswitz, F., J. O’Connell, 3rd, and R.M. Stroud, *Inhibitory complex of the transmembrane ammonia channel, AmtB, and the cytosolic regulatory protein, GlnK, at 1.96 A*. Proc Natl Acad Sci U S A, 2007. 104(1): p. 42–7.

27. Boeckstaens, M., et al., The TORC1 effector kinase Npr1 fine tunes the inherent activity of the Mep2 ammonium transport protein. Nat Commun, 2014. 5: p. 3101.

28. Loque, D., et al., A cytosolic trans-activation domain essential for ammonium uptake. Nature, 2007. 446(7132): p. 195–8.

29. Neuhauser, B., et al., Regulation of NH4+ transport by essential cross talk between AMT monomers through the carboxyl tails. Plant Physiol, 2007. 143(4): p. 1651–9.

30. Lanquar, V., et al., Feedback inhibition of ammonium uptake by a phospho-dependent allosteric mechanism in Arabidopsis. Plant Cell, 2009. 21(11): p. 3610–22.

31. Yuan, L., et al., Allosteric regulation of transport activity by heterotrimerization of Arabidopsis ammonium transporter complexes in vivo. Plant Cell, 2013. 25(3): p. 974–84.

32. Straub, T., U. Ludewig, and B. Neuhauser, The Kinase CIPK23 Inhibits Ammonium Transport in Arabidopsis thaliana. Plant Cell, 2017. 29(2): p. 409–422.

33. Beier, M.P., et al., Lack of ACTPK1, an STY kinase, enhances ammonium uptake and use, and promotes growth of rice seedlings under sufficient external ammonium. Plant J, 2018. 93(6): p. 992–1006.

34. Ma, L., et al., SALT OVERLY SENSITIVE2 and AMMONIUM TRANSPORTER1;1 contribute to plant salt tolerance by maintaining ammonium uptake. Plant Cell, 2025. 37(2).

35. Kirchhofer, A., et al., Modulation of protein properties in living cells using nanobodies. Nat Struct Mol Biol, 2010. 17(1): p. 133–8.

36. Hagn, F., M.L. Nasr, and G. Wagner, Assembly of phospholipid nanodiscs of controlled size for structural studies of membrane proteins by NMR. Nat Protoc, 2018. 13(1): p. 79–98.

37. Lei, J. and J. Frank, Automated acquisition of cryo-electron micrographs for single particle reconstruction on an FEI Tecnai electron microscope. J Struct Biol, 2005. 150(1): p. 69–80.

38. Scheres, S.H., RELION: implementation of a Bayesian approach to cryo-EM structure determination. J Struct Biol, 2012. 180(3): p. 519–30.

39. Zheng, S.Q., et al., MotionCor2: anisotropic correction of beam-induced motion for improved cryo-electron microscopy. Nat Methods, 2017. 14(4): p. 331–332.

40. Zhang, K., Gctf: Real-time CTF determination and correction. J Struct Biol, 2016. 193(1): p. 1–12.

41. Yang, J., et al., Improved protein structure prediction using predicted interresidue orientations. Proc Natl Acad Sci U S A, 2020. 117(3): p. 1496–1503.

42. Emsley, P. and K. Cowtan, Coot: model-building tools for molecular graphics. Acta Crystallogr D Biol Crystallogr, 2004. 60(Pt 12 Pt 1): p. 2126–32.

43. Adams, P.D., et al., PHENIX: a comprehensive Python-based system for macromolecular structure solution. Acta Crystallogr D Biol Crystallogr, 2010. 66(Pt 2): p. 213–221.

44. Davis, I.W., et al., MolProbity: all-atom contacts and structure validation for proteins and nucleic acids. Nucleic Acids Res, 2007. 35(Web Server issue): p. W375–83.

45. Pettersen, E.F., et al., UCSF ChimeraX: Structure visualization for researchers, educators, and developers. Protein Sci, 2021. 30(1): p. 70–82.

46. Marini, A.M., et al., A family of ammonium transporters in Saccharomyces cerevisiae. Mol Cell Biol, 1997. 17(8): p. 4282–93.

47. Hui, J., et al., The mycorrhiza-specific ammonium transporter ZmAMT3;1 mediates mycorrhiza-dependent nitrogen uptake in maize roots. Plant Cell, 2022. 34(10): p. 4066–4087.

48. Graff, L., et al., N-terminal cysteines affect oligomer stability of the allosterically regulated ammonium transporter LeAMT1;1. J Exp Bot, 2011. 62(4): p. 1361–73.

