## Supplementary Figure for "Structural basis for phosphorylation-induced inactivation of the rice ammonium transporter AMT1;1"

### Supplementary Figures S1-5

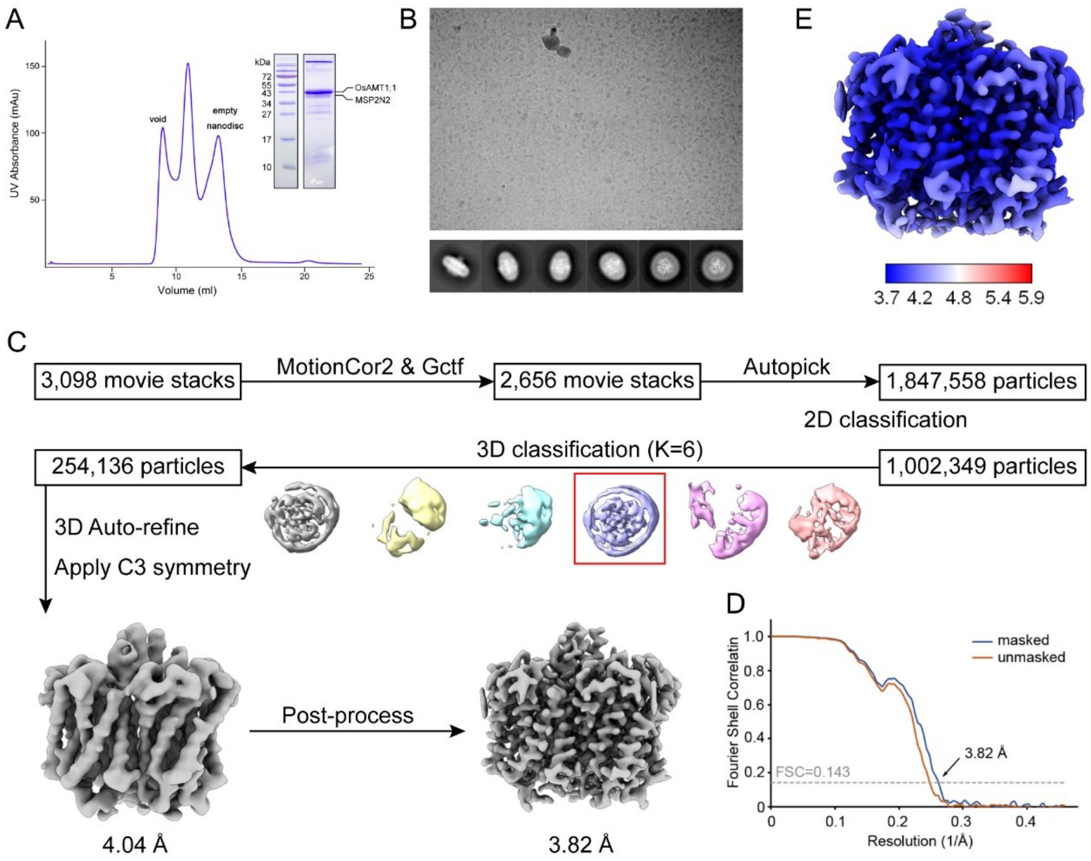


**Supplementary Figure S1. Cryo-EM analysis of OsAMT1;1.**

**A,** Gel filtration profile of a Superdex 200 column and Coomassie-blue-stained SDS-PAGE analysis of OsAMT1;1 reconstituted into MSP2N2. Independent experiments have been repeated at least three times with similar results. **B,** Representative micrograph and 2D class averages. **C,** Cryo-EM data analysis pipeline. **D,** Gold-standard Fourier shell correlation (FSC) curves. **E,** Local resolution estimation. The color represents the local resolution in Å.

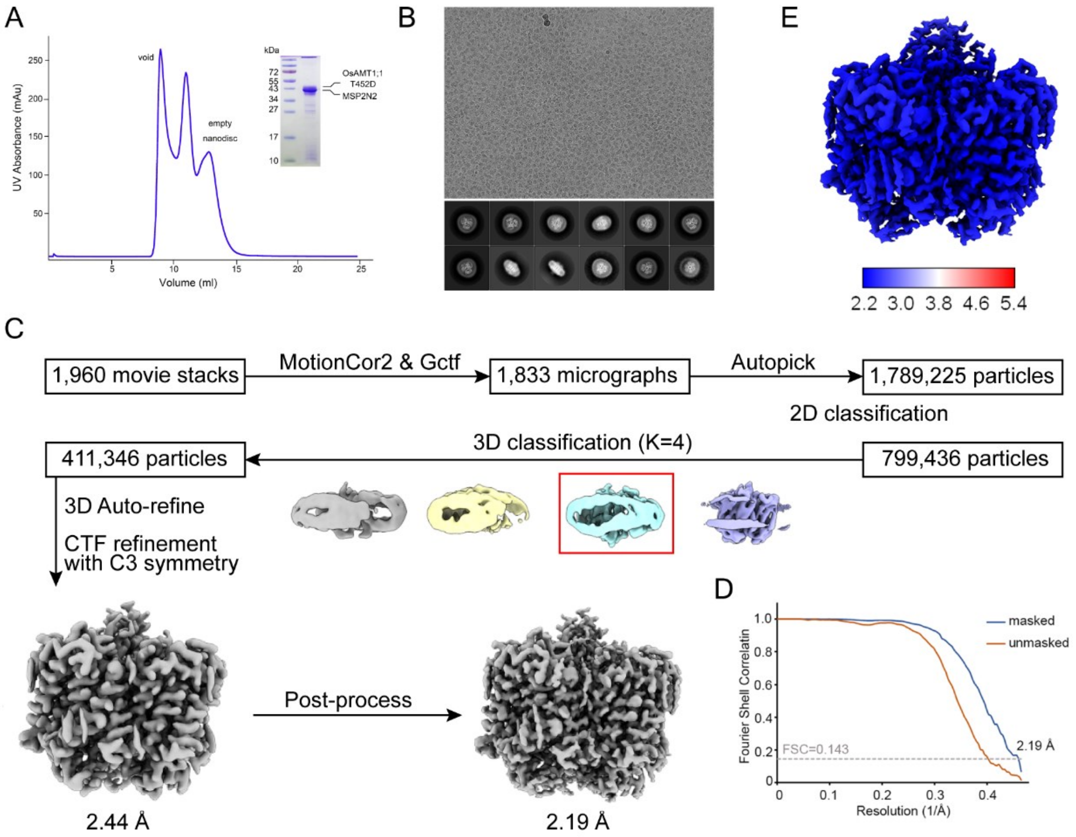


**Supplementary Figure S2. Cryo-EM analysis of OsAMT1;1 T452D.**

**A,** Gel filtration profile of a Superdex 200 column and Coomassie-blue-stained SDS-PAGE analysis of OsAMT1;1 T452D reconstituted into MSP2N2. Independent experiments have been repeated at least three times with similar results. **B,** Representative micrograph and 2D class averages. **C,** Cryo-EM data analysis pipeline. **D,** Gold-standard Fourier shell correlation (FSC) curves. **E,** Local resolution estimation. The color represents the local resolution in Å.

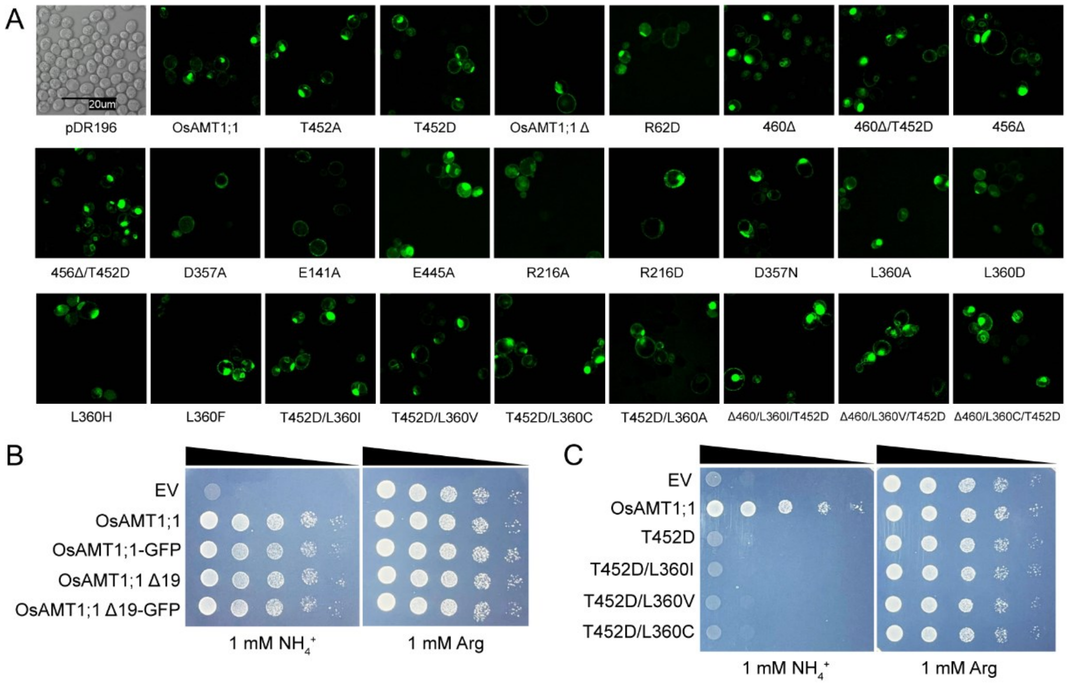


**Supplementary Figure S3. Functional complementation assays of OsAMT1;1 in yeast strain.**

**A,** Representative GFP fluorescence images of pDR196-OsAMT1;1-GFP wildtype, mutants and truncations in yeast strain. Yeast cells expressing empty vector serves as control. Scale bar, 20 μm. **B-C**, Complementation by pDR196-expressed OsAMT1;1 GFP-fusion (B) and phosphorylation site mutant T452D in combination with L360 mutants (C) supplemented with 1 mM arginine or 1 mM ammonium chloride. EV, empty vector.

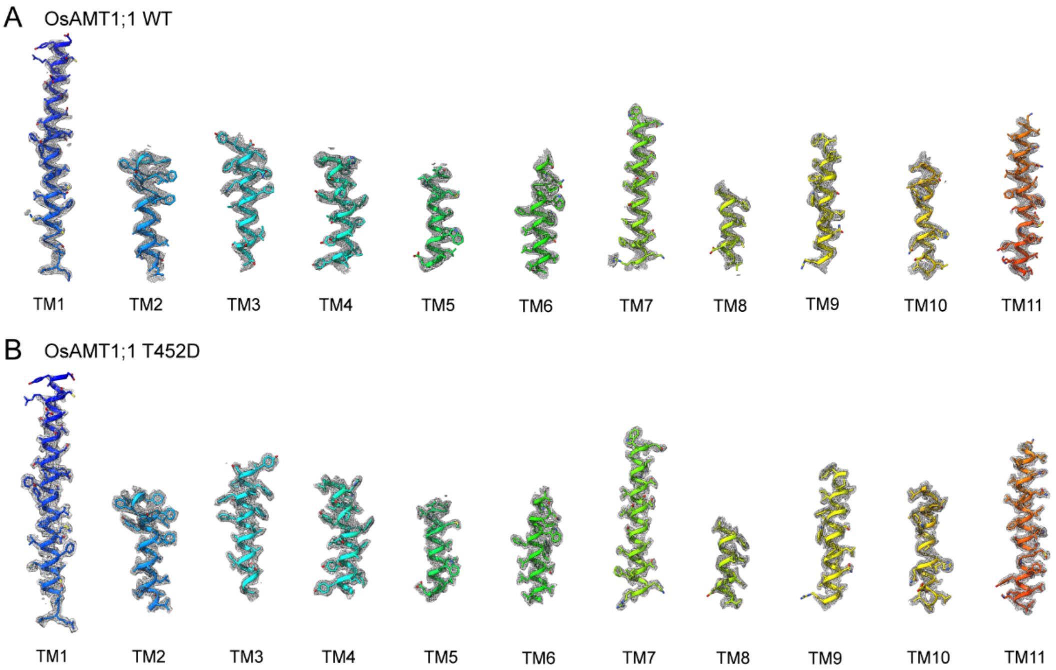


**Supplementary Figure S4. Representative densities of OsAMT1;1 WT and T452D.** Cryo-EM density of representative segments superimposed with the atomic model in OsAMT1;1 WT (A) and OsAMT1;1 T452D (B).

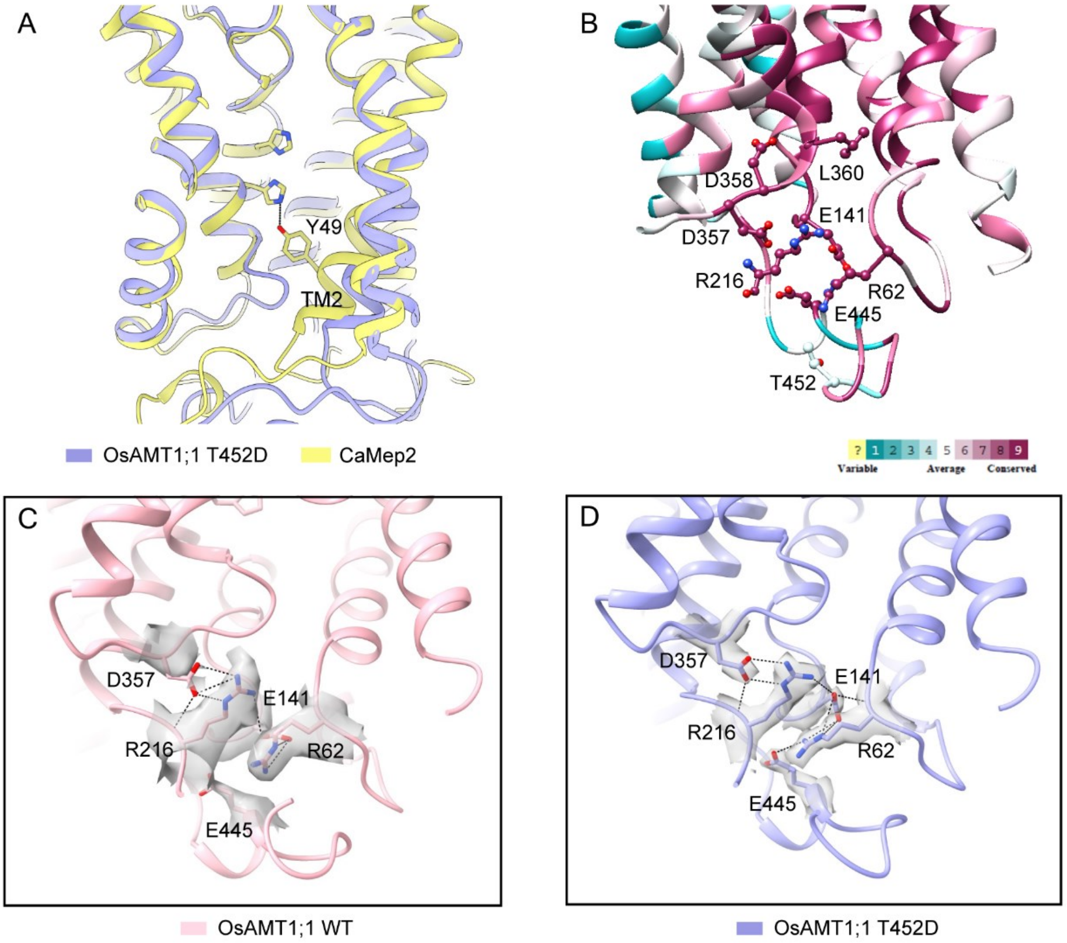


**Supplementary Figure S5. Salt-bridge network of OsAMT1;1 WT and T452D.**
**A,** Structural superposition of OsAMT1;1 T452D and CaMep2 (PDB 5AF1) monomers. The twin-His motif and residue Tyr49 in CaMep2 are shown as sticks. **B,** The sequence conservation analysis of OsAMT1;1. The residues involved in salt-bridge network are shown as sticks. **C-D,** Cryo-EM densities of salt-bridge network residues in OsAMT1;1 WT (C) and OsAMT1;1 T452D (D).

**Supplementary Table S1. Cryo-EM data collection, refinement and validation statistics.**

| | OsAMT1;1 (EMD-67958) (PDB 21SC) | OsAMT1;1 T452D (EMD-67959) (PDB 21SD) |
|---|---|---|
| **Data collection and processing** | | |
| Magnification | 81,000 | 81,000 |
| Voltage (kV) | 300 | 300 |
| Electron exposure ($e^-/Å^2$) | 50 | 50 |
| Defocus range (μm) | -1.2 to -2.2 | -1.3 to -1.8 |
| Pixel size (Å) | 1.09 | 1.077 |
| Symmetry imposed | C3 | C3 |
| Initial particle images (no.) | 1,847,558 | 1,789,225 |
| Final particle images (no.) | 254,136 | 411,346 |
| Map resolution (Å) | 3.82 | 2.19 |
| FSC threshold | 0.143 | 0.143 |
| Map resolution range (Å) | 3.82-5.90 | 2.19-5.40 |
| **Refinement** | | |
| Initial model used (PDB code) | trRosetta model | 21SC |
| Map sharpening *B* factor ($Å^2$) | -219.484 | -69.7946 |
| Model composition | | |
| Non-hydrogen atoms | 9825 | 9936 |
| Protein residues | 1305 | 1320 |
| Ligands | - | - |
| Water | - | - |
| *B* factors ($Å^2$) | | |
| Protein | 77.94 | 19.43 |
| Ligand | - | - |
| Water | - | - |
| R.m.s. deviations | | |
| Bond lengths (Å) | 0.005 | 0.005 |
| Bond angles (°) | 0.893 | 0.908 |
| Validation | | |
| MolProbity score | 1.57 | 1.37 |
| Clashscore | 3.85 | 3.00 |
| Poor rotamers (%) | 0.00 | 0.00 |
| Ramachandran plot | | |
| Favored (%) | 94.00 | 95.89 |
| Allowed (%) | 6.00 | 4.11 |
| Disallowed (%) | 0.00 | 0.00 |
